# Amino acids targeted based metabolomics study in non-segmental Vitiligo: a pilot study

**DOI:** 10.1101/2021.01.04.425209

**Authors:** Rezvan Marzabani, Hassan Rezadoost, Peyman Chopanian, Nikoo Mozafari, Mohieddin Jafari, Mehdi Mirzaie, Mehrdad Karimi

## Abstract

**Introduction:** Vitiligo is an asymptomatic disorder that results from the loss of pigments (melanin), causing skin or mucosal depigmentation and impairs beauty.

**Objective:** Due to the complexity of the pathogenesis of this disease and various theories including self-safety theory, oxidative stress, neurological theory and internal defects of melanocytes behind it, and finally, the vast role of amino acids in body metabolism and various activities of the body, amino acids targeted based metabolomics was set up to follow any fluctuation inside this disease.

**Methodology:** The study of amino acid profiles in plasma of people with non-segmental vitiligo using a liquid chromatography equipped with fluorescent detector was performed to find remarkable biomarkers for the diagnosis and evaluation of disease severity of patients with vitiligo. Twenty-two amino acids derivatized with *o*-phthalaldehyde (OPA) and fluorylmethyloxycarbonyl chloride (FMOC), were precisely determined. Next, the concentrations of these twenty-two amino acids and their corresponding molar ratios were calculated in 37 patients (including 18 females and 19 males) and corresponding 34 healthy individuals (18 females and 16 males). Using R programing, the data were completely analyzed between the two groups of patients and healthy to find suitable and reliable biomarkers.

**Results:** Interestingly, comparing the two groups, in the patient group, tyrosine, cysteine, the ratio of tyrosine to lysine and the ratio of cysteine to ornithine were increased while, arginine, lysine, ornithine and glycine ratios to cysteine have been decreased. These amino acids were selected for identification of patients with accuracy of detection of approximately 0.95 using the assessment of logistic regression.

**Conclusion:** These results indicate a disruption of the production of melanin, increased immune activity and oxidative stress, which are also involved in the effects of vitiligo. Therefore, these amino acids can be used as biomarker for the evaluation of risk, prevention of complications in individuals at risk and monitoring of treatment process.

## Introduction

Vitiligo is a common chronic skin disorder in which pigment-producing cells, or melanocytes are getting in trouble that can result in varying patterns and degrees of skin depigmentation. Patients are characterized by loss of epidermal melanocytes and progressive depigmentation. It is appeared in two main types, non-segmental (generalized) or segmental (Armstrong, 2011; Sahoo et al., 2017).

Regardless of much research, the etiology of vitiligo and the reasons of melanocyte death are still unclear (Singh et al., 2016). A complex immune, genetic, environmental, and biochemical causes are behind Vitiligo and the exact molecular mechanisms of its development and progression is not clear (Liang et al., 2019; Sahoo et al., 2017; Singh et al., 2016). Although several vitiligo susceptibility loci identified by genome-wide association studies were reported, but study examining monozygotic twins reported a vitiligo concordance rate of 23%, suggesting a strong environmental contribution to the pathogenesis (Singh et al., 2016). Zheleva et al. (Zheleva et al., 2018), in their work, revealed oxidative stress is a triggering event in the melanocytic destruction and is probably involved in the etiopathogenesis of vitiligo disease. Oxidative stress biomarkers could be finding in the skin and blood of vitiligo patients. Hamidizadeh et al. (Hamidizadeh et al., 2020), in their study compered hopelessness, anxiety, depression and general health of vitiligo patients in comparison with normal controls and confirmed that anxiety and hopelessness levels were significantly higher in vitiligo patients than those who are in healthy controls.

It is northly to know, vitiligo worldwide prevalence is in the range of 0.5% to 2% (Ding et al., 2014). But, one the main problems accompanied with vitiligo is its psychological aspect that is experienced by many patients around the globe (Grimes and Miller, 2018). Next to social or psychological distress, people with vitiligo may be at increased risk of sunburn, skin cancer, eye problems, such as inflammation of the iris (iritis) and hearing loss (Jakku et al., 2019). There are many both conventional and unconventional therapies for vitiligo. They are including L-phenylalanine, PGE2 and antioxidant agents, Alpha Lipoic Acid, Flavonoids, Glutathione (GSH), Fluorouracil, L-DOPA, Levamisole, L-Phenylalanine, Melagenine, Omega-3 polyunsatured fatty acids, cream cointaning Pseudocatalase, Resveratrol Soybeans, Metals such as zinc, Minoxidil (Gianfaldoni et al., 2018)

Although, the pathophysiology of vitiligo is complex, the studies revealing vitiligo cells have unique lipid and metabolite profiles (Sahoo et al., 2017). This led to the question of which factors been associated with vitiligo activity in skin and blood. These biomarkers allow an early and accurate determination of treatment response and the progression of the disease. Up to now some biomarkers is recommended for Vitiligo. Several markers which are received linked to vitiligo and associated with disease activity. Besides providing insights into the driving mechanisms of vitiligo, these findings could reveal potential biomarkers. Although genomic analyses have been performed to investigate the pathogenesis of vitiligo, but the role of small molecules and serum proteins in vitiligo remains unknown. providing insights into the driving mechanisms of vitiligo, these findings could reveal potential biomarkers. Metabolomics is a powerful and promising analytical tool that allows assessment of global low-molecular-weight metabolites in biological systems. It has a great potential for identifying useful biomarkers for early diagnosis, prognosis and assessment of therapeutic interventions in clinical practice (Liang et al., 2019; Speeckaert et al., 2017).

Despite the current evidence of the effects of metabolic system on immune system and oxidative stress as two important factors in the development of vitiligo, it seems necessary to more investigation of metabolite fluctuation in this disease. We were keen to establish whether levels of important substrates such as amino acids as the most important primary metabolites were altered in vitiligo cells. This might therefore contribute to the vitiligo phenotype in melanocytes. Then, the aim of this study was to investigate a comprehensive profile of amino acids in plasma of people with vitiligo in comparison with healthy people to find a fast-determinable biomarker. For this a liquid chromatography equipped with fluorescent detector was applied.

## Material and methods

### Patient samples

After receiving ethical approval (The study protocol was approved by the ethics committee of our institution. Also, informed consent was obtained voluntarily from each participant at the time of enrollment) from the Shahid Beheshti University of Medical, all participants signed written informed consent. Table 1 is demonstrating the complete characterization of the case studies. In summary 37 cases with vitiligo and 33 healthy ones attended to the dermatology clinic of Shohadaye Tajrish Educational Hospital. The diagnosis of vitiligo was based on the characteristic loss of skin pigmentation and the examination under Wood’s lamp.

**Table 1.**
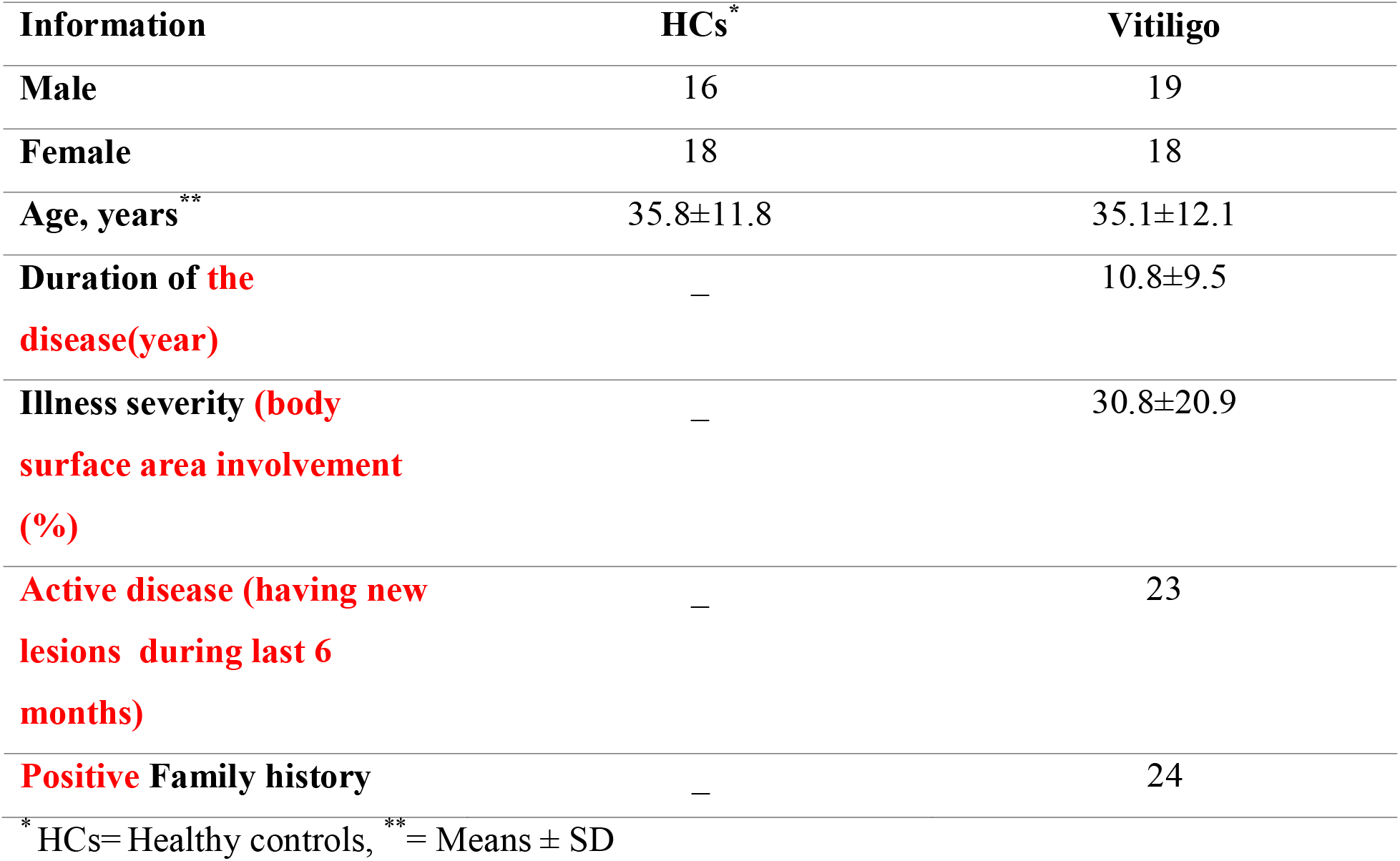
Demographics of the study cohort

Blood samples were entered in the tube vacutainers 10 mL containing 0.15 K_2_EDTA (to prevent clotting) and were centrifugal at 4000 rpm at 4 °c for 20 minutes. Supernatant was isolated and reserved for HPLC-FD analysis at −80 °C.

### Amino acid Analysis

In order to prepare the samples for analysis, the samples are transferred from −80 ° C refrigerator and placed in the ice to be melted. To 50 μL of sample 20 norleucine (500 μM) and then 200 μL of methanol kept at −20 °C and all are mixed for five seconds. To completely deproteination, they are kept at −20 °C for 2 hours. At the next stage, The samples are centrifuged at 13000 rom for twelve minutes at 4 °C. The supernatant is completely transfered to Heidolph rotary evaporator and dried in vacu. These samples could be reserved at 4 °C for four weeks.

For HPLC analysis, previously dried samples were dissolved in 100 μl of water (containing 0.01 formic acid) with help of ultrasonic device for 5 minutes. To 10 μl of each sample 10 μL OPA (for derivatization of primary amino acids) and one minute late 10 μL FMOC (for secondary amino acid derivatization) 20 μL of this sample are injected HPLC column (Fekkes, 2012; Wu et al., 2016)

For the HPLC-DAD method, a Knauer system (WellChrom, Germany) equipping with a K-1001 pump, a K-2800 fast scanning UV detector with simultaneous detection at four wavelengths, an autosampler S3900 (Midas), a K-5004 analytical degasser, and a 2301Rheodyneinjector with a 20 μL loop was used. HPLC separation was achieved using a Eurospher C18 column (4.6 mm × 250 mm, 5 μm), with a gradient elution program at a flow rate of 1.0 ml min-1. The mobile phase was composed of A (acetonitrile + 0.05% three flouro acetic acid, v/v) and B (0.05% aqueous trifluoroacetic acid, v/v). The following gradient was applied: 0–10 min, isocratic gradient 70% B; 10–30 min, linear gradient 70–40% B; 30–40 min, linear 40–20% B; 40–50 min, linear 20–0% B; 50–65 min, linear 0–70% B; 65–75 min, isocratic gradient 70% B. The UV absorbance was monitored at 335 nm. All injection volumes of sample and standard solutions were 20 μl. The chromatographic peaks of the sample solution were identified by spiking and comparing their retention times and UV spectra with those of reference standards. Quantitative analysis was carried out by integration of the peak using the external standard method. Identification of amino acids were conducted using fluorescence at 337 nm and 470 nm for adsorption and excitation respectively for primary amino acids and while detectors and 262 wavelengths (for second amino acids) and 338 nm (for first-type amino acids) related to PDA detectors. To check the accuracy of the procedure, five plasma samples related to individuals The patients were analyzed that RSD every 22 amino acids less than 7 were obtained (Amorini et al., 2017; Douglas, 2003; Wu et al., 2016).

### Statistical methods

For statisticalAnalysis, we used Metaboanalyst 4.0. Before the analysis, we applied The data conversion and the Mean Center scale and finally the data with normal normalize quantile (the data were analyzed by Shapiro-Wilk test in software R and the data was not normal for some amino acid¬). To compare between study groups by R software, we used Mann-Whitney U test with the FDR correction (benjamini Hochberg) 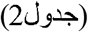. In addition to comparing two groups of patients and healthy patients, the relationship between severity of disease, disease activity, family history, and duration of each amino acid was usedto evaluate Mann Whitney U tests with FDR correction (benjamini Hochberg) and the average prediction score (Random forest) (table 5, image 2). In examining the trend of difference (variation Figure 6) in the amount of metabolites, the sample was used in two groups and clustering of partial separator (PLS-DA) method(Figure 7). In addition, to compare and investigate the correlation between two to two amino acids at the same time in all participants, the correlation matrix was plotted with a significant difference asaheatmap(Form5). It was also plotted to investigate the relationship between each amino acid and the participants of the heat map (Figure 8). Also, in order to investigate the effect of each amino acid and their ratios (20 superior ratio based on Pvalues) We selected as biomarker in the expression of the probability of the cause or severity of the disease, we used logistic regression, the results of the sensitivity and specificity of the test and the result of the system performance curve (ROC) Multiple queries (image 6).

## Results

Totally determined 22 amino acids were determined in the studied samples. Table S1. is demonstrating the absolute concentration for determination of twenty-two amino acids in participant group (34 healthy cases and 37 vitiligo cases). First, we performed principal component analysis with all samples, which showed that samples were well clustered in two completely separated clusters (Figure. 1).

**Figure 1.**
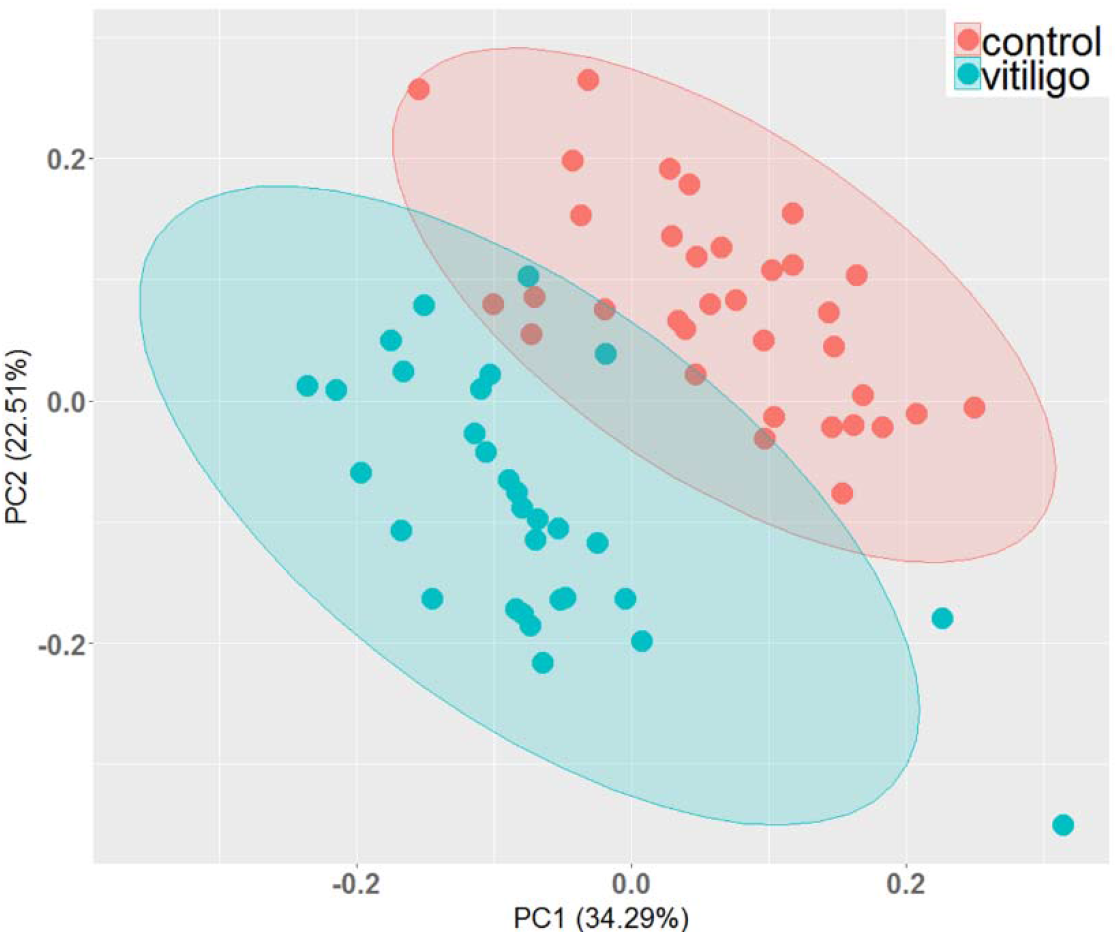
PCA analysis shows the homogeneity of data obtained by HPLC-FLD. Samples are completely grouped to tow separated cluster. PC1 and PC2 are covered 34% and 22% all data obtained.

Next, Amino acid distribution was evaluated by Shapiro test. Also, *t*-test was used to show amino acids differences in concentration between vitiligo and healthy samples. Adjusted *p*-values calculated by Benjamini Hochberg methods. Figure 2(a). is demonstrating volcano graph in which horizontal and vertical axes are corresponding to log2 fold change of sample concentrations and −log10 adjusted *p*-values respectively. As illustrated in figure 2(c-d), there is a significant increasing in Cys, Pro and Glu, while Lys, Arg, Orn, His and Gly are decreased in vitiligo patients. Figure 2(b). is showing Gini error reduction diagram (average accuracy reduction, average prediction score) obtained from Random Forrest algorithm with tree number of 500. The green dots in vitiligo have increased and the red dots in vitiligo have decreased.

**Figure 2.**
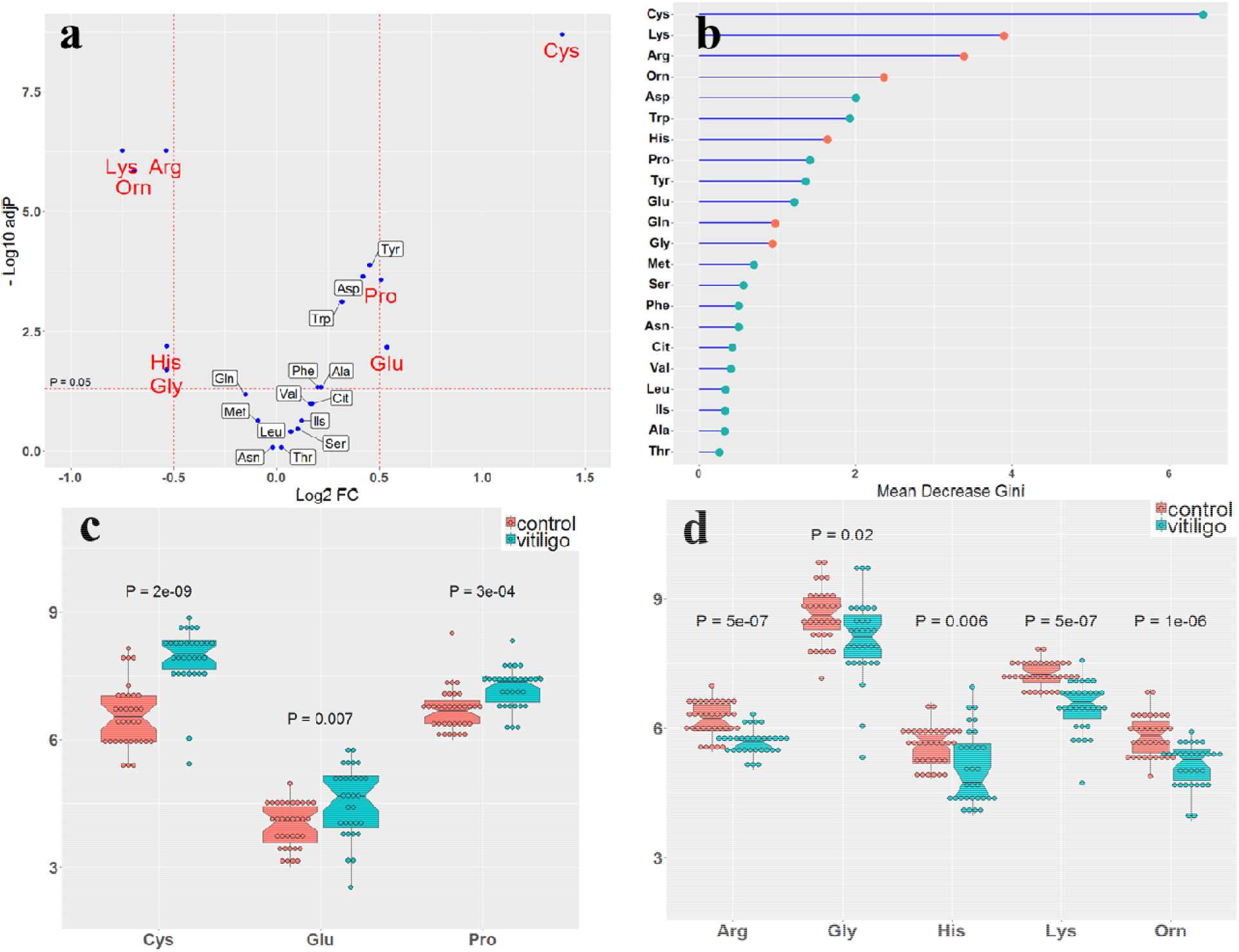
**(a)** Volcano graph related to amino acids concentration change in the studied Vitiligo samples, (b) Gini error reduction diagram with tree number of 500, (c-d) Box-plot for amino acids fluctuation in both healthy and vitiligo samples. The red boxes show the metabolic concentration values in healthy individuals (control), while the blue ones show the metabolic concentration values in the sick individuals (vitiligo). The adjust *p*-values for each metabolite are mentioned in the figure.

To show the specificity and sensitivity of the studied biomarkers, ROC graph was used. Also, an individual ROC curve was plotted for amino acids with highest changes (Figure 3(b). Interestingly, Cys and Lys showed the maximum of area under curve (AUC) up to 0.91. For these two amino acids a logistic regression was done and its corresponding ROC diagram was drawn. Positive/negative coefficient is implying to the role of each of the selected amino acids in Increasing or decreasing the risk of vitiligo. Next, based on random forest method a confusion matrix developed in which two group of our study are completely classified (Figure 3).

**Figure 3.**
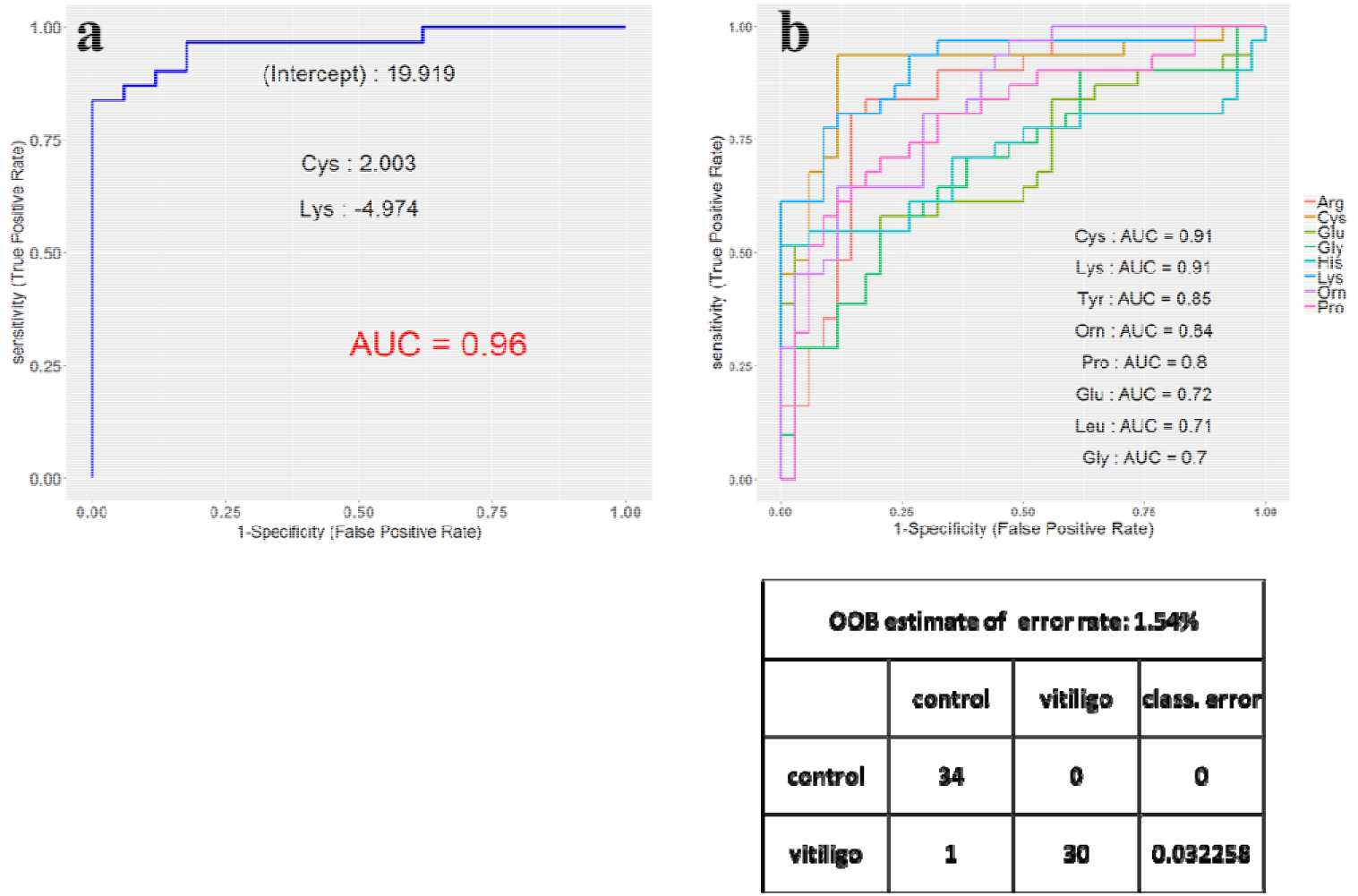
**(a)** ROC curve to show the sensitivity and specificity of the studied amino acids (Cys, Lys, Tyr, Orn, Pro, Glu, Leu, and Gly), (b) Selected ROC curve for Cys and with the highest variation, (c) confusion matrix, based on random forest is completely

Following the question on the variation of amino acids concentration inside Vitiligo cases with two category, more than 25% and less than 25%, Glu found to be a reliable biomarker. Its concentration (Log2 FC< −0.5) is significantly decreased in the patient showing more than 25% (Figure 4(a-b)).

**Figure 4.**
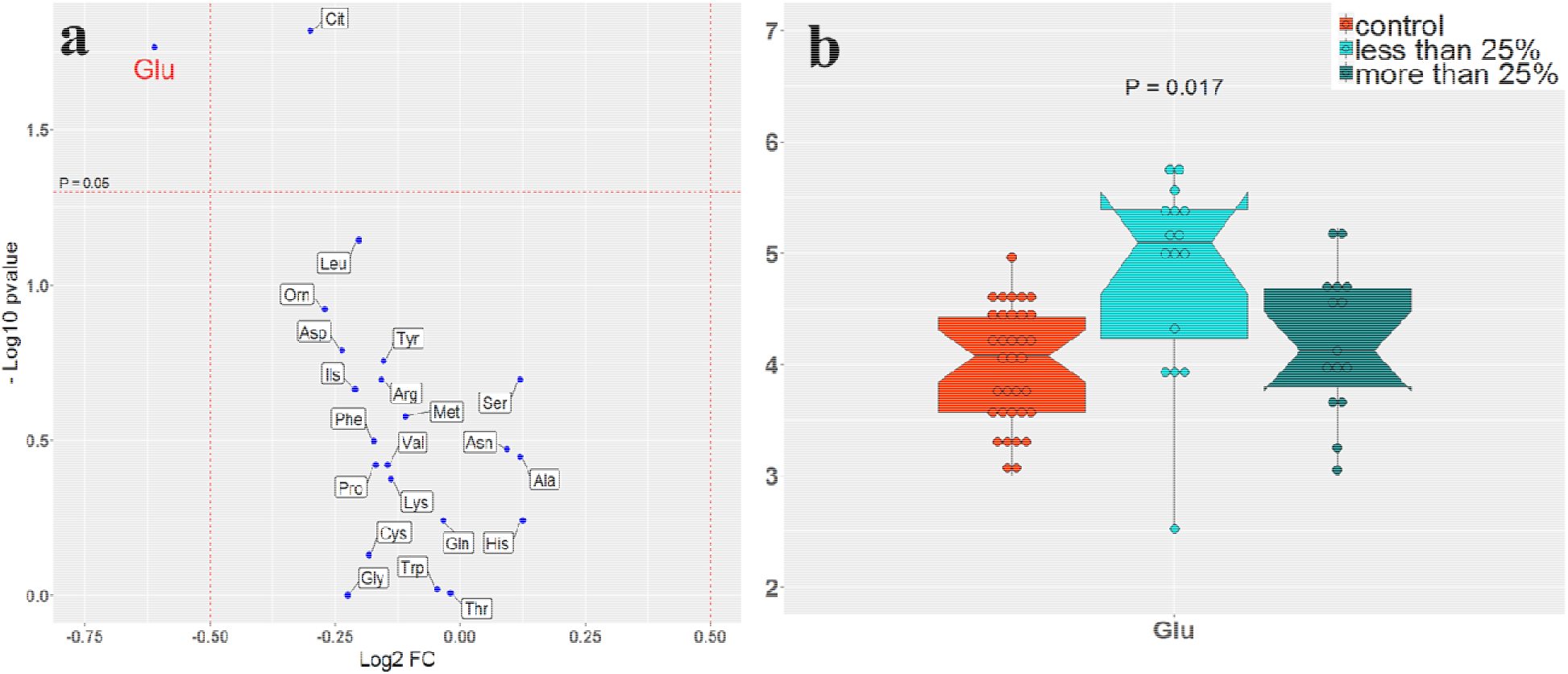
Volcano diagram related to patients with more and less 25% of Vitiligo. (a) Glu is classifying the cases according to Vitiligo severity, (b) Glu is decreased in Patina t with more than 25% of Vitiligo.

As the ratio of biomarkers especially amino acids would be a reliable sign of disease, volcano diagram for different ratio of amino acids in the Vitiligo samples are prepared. According to figure 5, ratios including Cys/Orn, Gly/Cys, and…are significantly group the cases of the study.

**Figure 5.**
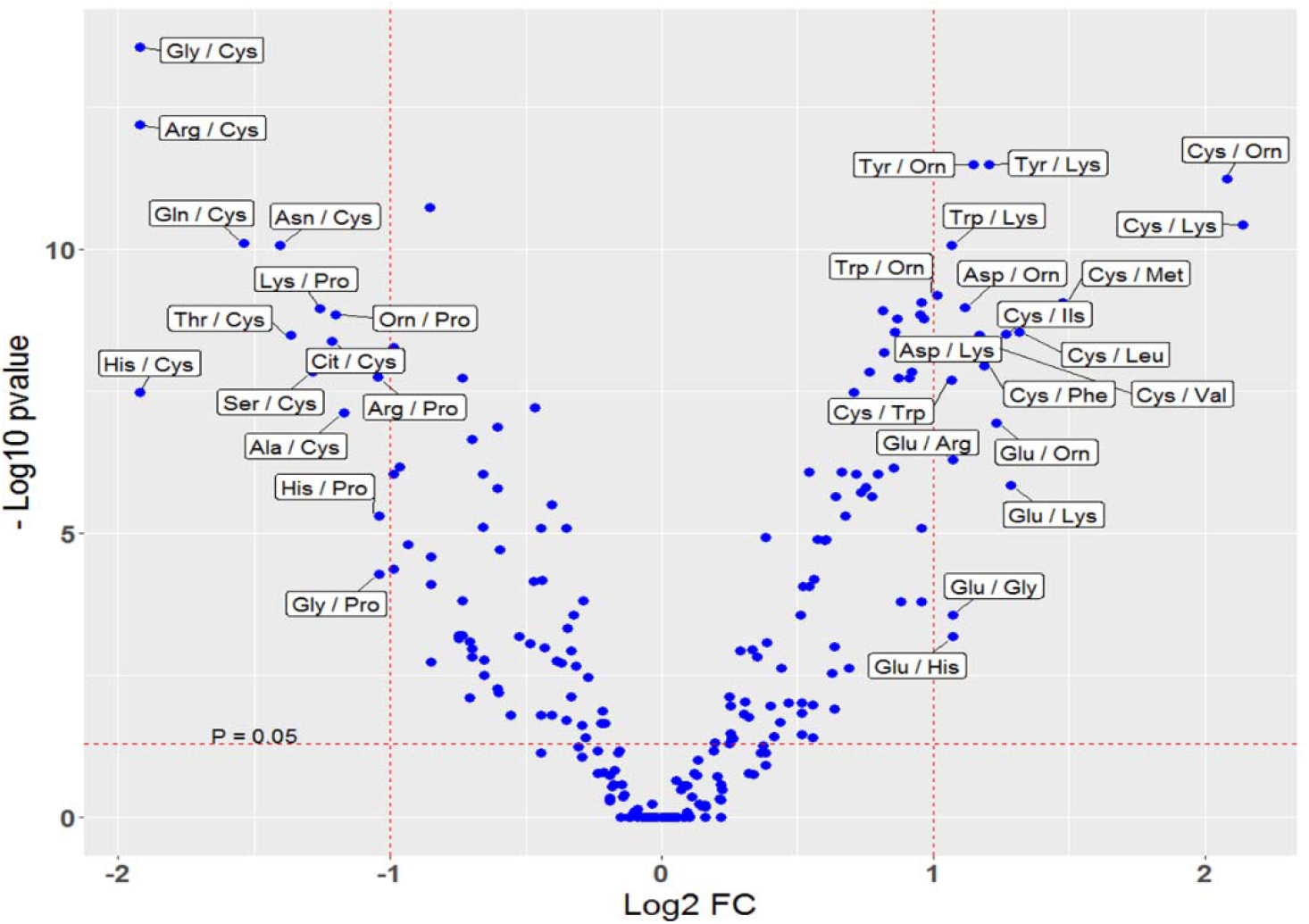
Volcano diagram related to amino acid variation between the studied cases.

**Figure 6.**
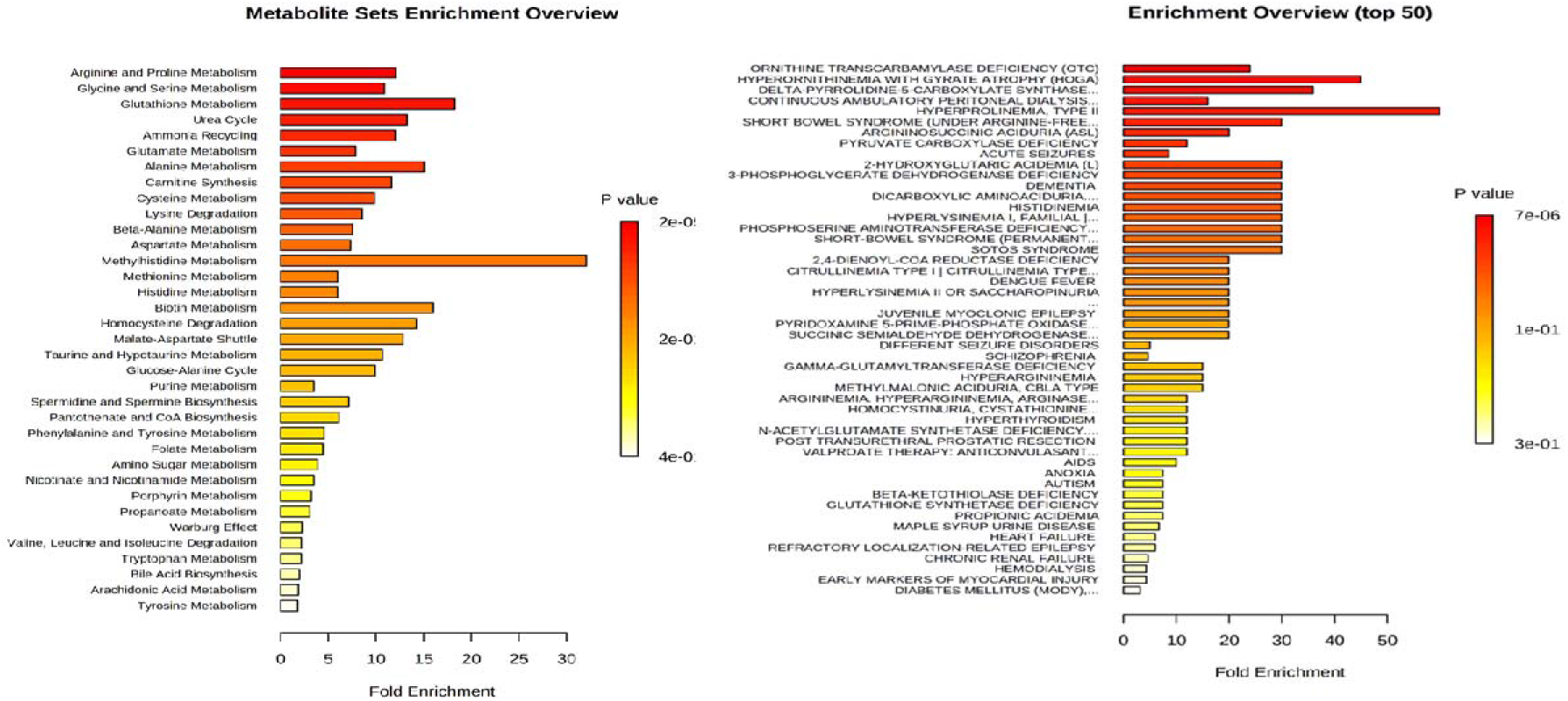
Pathway-associated metabolite and disease-associated metabolite analyses

**Figure 7.**
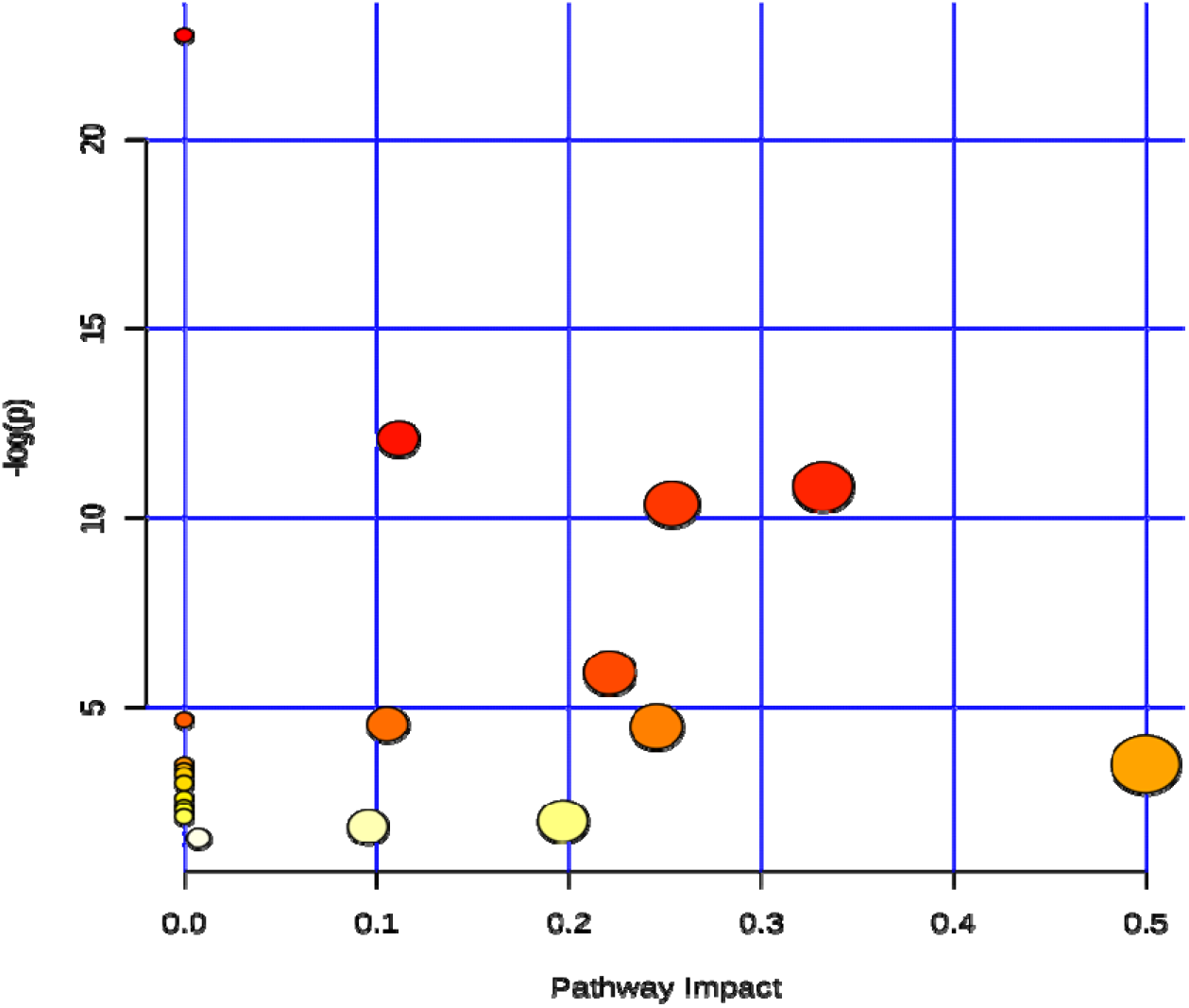
Specify pathway analysis algorithms: Over Representation Analysis: Hypergeometric Test Pathway Topology Analysis: Relative-betweeness Centrality

### Metabolic Pathway Analysis

Metabolite set enrichment analysis (MSEA) was used to explore the metabolites highly enriched and associated with possible metabolic pathways. Pathway-associated metabolite and disease-associated metabolite analyses were performed shows the majority of the metabolic pathways that are significantly altered in Vitiligo cases.

Using pathway associated metabolite sets with enrichment analysis, the main pathways affected were detected. Pathway impact as checked by Metaboanalyst??? has shown that about 35 pathways differ between Vitiligo and healthy samples, of which the first 12 pathways are very significant. following metabolites and metabolic cycles are found to be changed in Vitiligo cases: Arginine and proline metabolism, glycine and serine metabolism, glutathione metabolism, urea cycle, ammonia recycling, glutamate metabolism, alanine metabolism, carnitine synthesis, cycteine metabolism, lysine degradation, beta-alanine metabolism, aspartate metabolism, and methyl histidine metabolism. These are pathways and metabolic cycles, which differed significantly between VCs and HCs.

On the other hand, disease-associated metabolite sets compared between VCs and HCs. Ornithine transcarbamylase deficiency (OTC), Hyperornithinemia with gyrate atrophy (HOGA), Delta-pyrrolide-5-carboxylate synthase, continuous ambulatory peritoneal dialysis, hyperprolinemia-type II, short bwel syndrome (under arginine -free), Argininosuccinic aciduria (ASL), acute seizures, 2-hydroxyglutaric acidemia, 3-phosphoglycerate dehydrogenase deficiency dementia, dicarboxylic aminoaciduria, histinemia, hyperlysinemia I-Family I, phosphoserine aminotransferase deficiency, short-bwel syndrome, and SOTOS syndrome are the most disease-associated metbaile we found here.

## Discussion

To best of our knowledge, there are few studied on the role of amino acids in vitiligo. They are focus only on the one or two amino acids and their metabolites which are associated with the production pathway of melanin (phenylalanine, tyrosine and glucosamine, trimethylamine, cysteine, homocysteine and thiol). However, no studies have been conducted to investigate the profile of free amino acids, to investigate changes in those and metabolic pathways of vitiligo. Amino acids play an important role in detoxification and immune responses through regulating the activation of T lymphocytes, B lymphocytes, natural killer cells, and macrophages (1), cellular redox state, gene expression, and lymphocyte proliferation (2), and the production of antibodies, cytokines, and other toxic compounds for the cell (3).

In most of the cell types, arginine is produced from citrulline as a precursor and is involved in regulating the activity of the immune system by producing nitric oxide. Proline and glutamate synthesize ornithine by producing pyrroline-5-carboxylate (P5C). In addition, it is catabolized by proline oxidase in different organs to produce hydrogen peroxide and P5C. By converting P5C into proline, a reduction occurs in the ratio of NADP^+^ to P5C reductase-dependent NADPH. The proline-P5C cycle regulates the cellular redox state and cell proliferation. In addition, ornithine is converted into citrulline and regenerates arginine using aspartate. Given the metabolic pathways, in which arginine is involved, each of its products has a specific function, including ornithine, as a product of arginine, proline, and glutamate, which contributes to the production of glutamate, glutamine, and polyamines, and mitochondrial integrity, polyamines, as the products of arginine and methionine, affect gene expression, DNA and protein production, ion channel activity, cell death, antioxidants, cellular activity, proliferation and differentiation of lymphocytes, and creatine, as a product of arginine, methionine, and glycine, has antioxidant, antiviral, and antitumor activity. Therefore, concomitant decrease in arginine and ornithine and increase in proline may indicate impaired arginine and proline metabolism and urea cycle. As a result, there is a disruption in the response to oxidative stress and cell damage.

There are several serine-pathways involving one-carbon metabolism, one of which is glycine synthesis. Glycine is involved in synthesizing many important physiological molecules, including purine nucleotides, glutathione, and Heme (a cofactor containing an iron atom). In addition, glycine itself is a potent antioxidant scavenging free radical. Therefore, glycine is essential for the proliferation and antioxidative defense of leukocytes, and is an antiinflammatory, immunomodulatory, and cytoprotective agent, the reduction of which indicates impaired glycine/serine and glutathione metabolism, which, in turn, disrupts cellular immunity and response to oxidative stress.

Ammonia is considered as an important source of nitrogen and a by-product of cellular metabolism. In addition, it is absorbed through reducing amine synthesis catalyzed by glutamine synthetase and glutamate dehydrogenase, the secondary reactions of which enable other amino acids such as glutamate, proline, and aspartate to obtain this nitrogen directly. Glutamate regulates the expression of nitric oxide synthases (iNOS) in specific tissues and is indirectly involved in regulating the animal immune system. Aspartate, acting as a precursor for nucleotide synthesis, contributes to various metabolic pathways and is important for lymphocyte proliferation. Further, it is necessary for regenerating arginine produced from citrulline in active macrophages and maintaining the intracellular concentration of arginine to sustain NO level in response to immune challenges. Glutamate and aspartate play stimulating roles in the central and peripheral nervous systems, affecting ionotropic and metabotropic receptors (peptide or polypeptide hormone receptors and neurotransmitters on the plasma membrane which play an important role in the immune system). They transport the reducing agents across the mitochondrial membrane, thereby regulating glycolysis and cellular redox state through the malate/aspartate shuttle. In addition, alanine, as a major substrate for hepatic glucose synthesis, is a significant energy substrate for leukocytes, thereby affecting immune function. /?-alanine is the only non-essential beta amino acid which occurs naturally and is formed by various metabolic organs. Additionally, they are involved in producing glutamate, aspartate, glutamine, and glycine in a part of their metabolic pathways. Aspartate and glutamate, along with glutamine, are the main source of energy for enterocytes (intestinal epithelial cells). The results showed that glutamate increased among the patient group compared to the control group, indicating increased immune system activity and impaired cellular redox state.

Methionine is converted into homocysteine (used as a source of sulfur) in the course of its metabolism and cysteine is produced after homocysteine binds to serine and an intermediary cystathionine is formed. Some studies examined homocysteine and thiols in vitiligo patients and found an increase in homocysteine due to essential cofactors and folate for the activity of methionine synthetase and, consequently a decrease in its activity and the methionine reproduction cycle, which led to an increase in cysteine. Tyrosine is converted into dopaquinone, a highly intermediary metabolite, by tyrosinase which is important for regulating melanogenesis.

Dopaquinone reacts rapidly with cysteine as it increases to get involved in the production of pheomelanin, which is considered as a common type of melanin pigment found in the hair and skin, the color of which changes from yellow to red as its concentration increases. When the cysteine level does not decrease, the reaction does not lead to the production of eumelanin pigment, the increased concentration of which changes the color from light brown to black [1]. By increasing thiol levels, the production of melanin is impaired. In addition, the dynamic thiol/disulfide homeostasis regulates the storage of antioxidants, detoxification, apoptosis, and many signal mechanisms including cell division and growth. The results indicated that an increase in cysteine and ratio of cysteine to ornithine and a decrease in the ratio of glycine, arginine, ornithine, and lysine to cysteine in the patient group. Thus, impaired cysteine metabolism disrupts pigment production, increases the activity of the immune system, and counteracts the effects of oxidative stress due to the deficiency in the production of antioxidant compounds such as taurine and glutathione, which result in damaging melanocytes and decreasing pigments.

Lysine, which reduced in the group of vitiligo patients, has multiple catabolic pathways, the main one of which is in the liver, where saccharopine, glutamate, alpha-aminoadipate 6-semialdehyde, and acetyl-CoA are produced [2]. In the human body, carnitine, involved in fatty acid metabolism, is biosynthesized using amino acids lysine and methionine. Carnitine and its esters help reduce oxidative stress [3]. In addition, dietary or extracellular lysine can modulate the entry of arginine into leukocytes and the production of NO by iNOs through sharing the like transport systems with arginine.

Histidine is converted into urocanic acid through one of its metabolic pathways by enzymatic catalysis of histamine ammonia-lyase. UCA is a unique photoreceptor and cis-UCA is converted into trans-urocanic acid (trans-UCS) by absorbing ultraviolet (UV) radiation from the sun, which controls the activity of the immune system against the UV radiation from the sun. Increased or decreased histidine level from the normal state disrupts the function of the skin immune system [4, 5]. Decreased histidine in the patients triggers the activity of the immune system in response to the existing stimuli, making their skin cells more vulnerable to UV radiation than the normal state.

3-methylhistidine is formed by the posttranslational methylation of histidine residues from major myofibrillar proteins (actin, and myosin). In humans, it is associated with a variety of diseases including type 2 diabetes, eosinophilic esophagitis, and kidney disease. In addition, 3-methylhistidine is associated with the metabolic disorder of propionic acidemia. Measuring 3-methylhistidine provides an indicator of the rate, at which muscle protein breaks down. It is also a biological marker for meat intake, muscle protein breakdown, and intestinal proteins.

The clinical features of vitiligo are classified in different ways, one of which is based on the extent of the spots on the body surface. In the patients who were divided into two groups, with limited extent of spots (less than 25%) and large spots (greater than 25%), glutamate decreased by increasing spot extent. Due to the role of glutamate in regulating the protein synthesis and breakdown in the cell and cell cycle, its lower level in these people can indicate impaired cell metabolism and increased cell death, resulting in increased complications in people with more severe disease.

Given these cases, the meta-analytic recommendations for the diseases associated with impaired pathways are better understood such as ornithine transcarbamylase (OTC) deficiency (an inherited disorder that causes ammonia to accumulate in the blood due to deficiency in the transcarbamylase), hyperornithinemia with gyrate atrophy (HOGA) (an inherited disorder characterized by progressive vision loss). Disruption of ornithine aminotransferase production helps convert ornithine into another molecule, called P5C. P5C can be converted into amino acids (glutamate and proline), delta-pyrrolide-5-carboxylate synthase (difficulty in degrading proline to P5C), continuous ambulatory peritoneal dialysis (difficulty in excreting all urea and ammonia and, therefore, the need for dialysis), hyperprolinemia-type II (problems with proline degradation increase proline and P5C), short bowel syndrome (under arginine-free) (the small intestine is required for arginine synthesis). Therefore, limited access to essential amino acids in the patients with SBS leads to a defect in the intermediates of the urea cycle, ornithine, citrulline, and arginine, as well as a reduction in these amino acids, which may lead to hyperammonemia, argininosuccinic aciduria (ASL), as a urea cycle disorder which causes ammonia to accumulate in the blood. Other suggested disorders are all inherited diseases which cause complications and metabolic disorders.

Based on the results obtained from the review of data and results of previous studies in this area, it is observed that reduced melanin production due to increased cysteine in the patients as well as autoimmunity, and oxidative stress (increased glutamic acid and proline and decreased arginine, glycine, lysine, histidine, and ornithine in patients) simultaneously can damaging melanocytes, result in vitiliginous lesions on the skin surface of patients. Thus, examining the proposed biomarkers may be helpful in early diagnose of at risk patients, in addition considering the changes in glutamic acid levels as biomarkers can be useful for determining the prognosis of the disease. Also Understanding the role of these biomarkers in vitiligo can provide the scientific basis for the development of novel therapeutic approaches in this disease.

## Conclusion

