## Supplementary figures and images for "Amino acids targeted based metabolomics study in non-segmental Vitiligo: a pilot study" (from another version of this preprint: version 1, 2021-01-05)

### Supplement 2

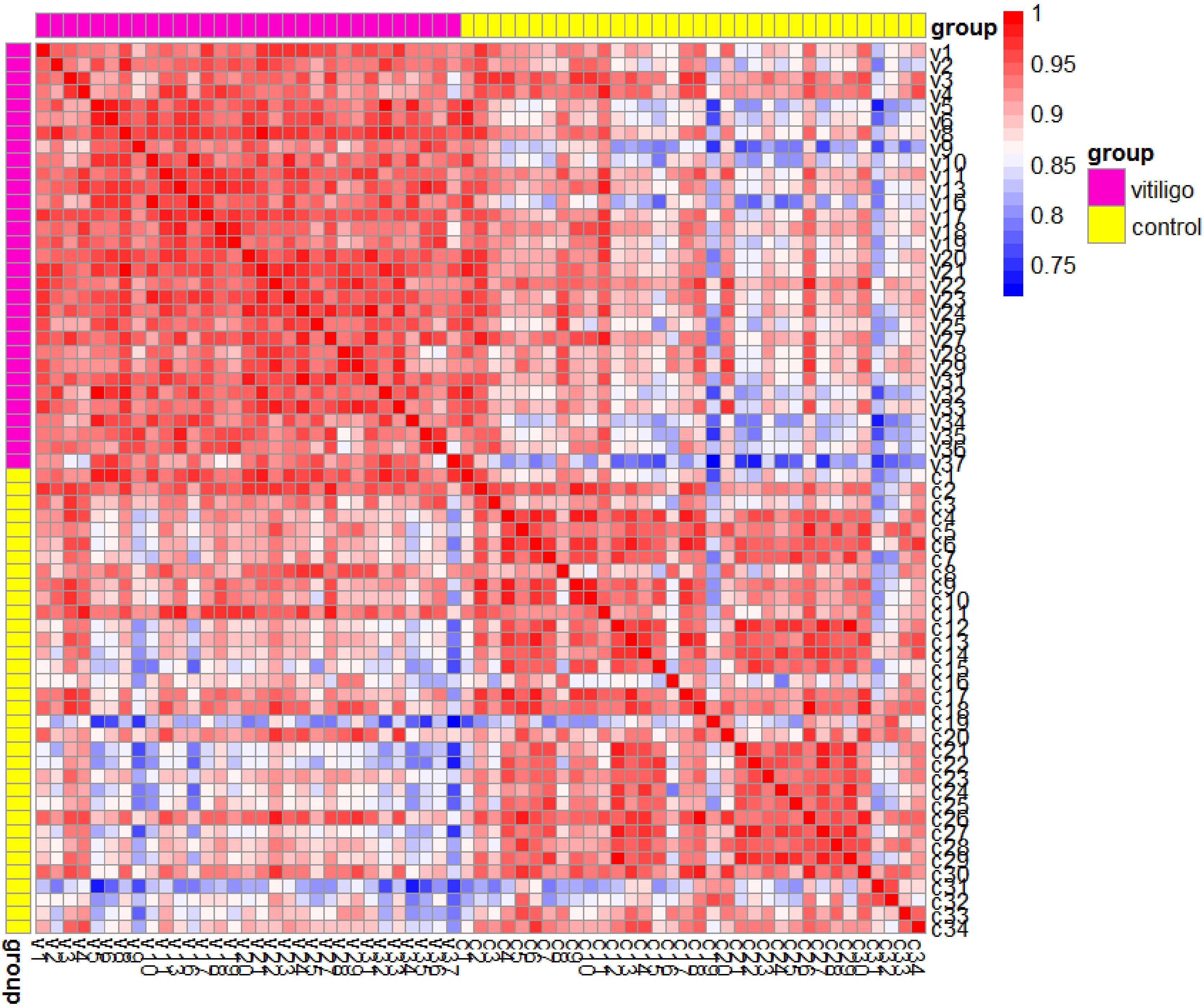

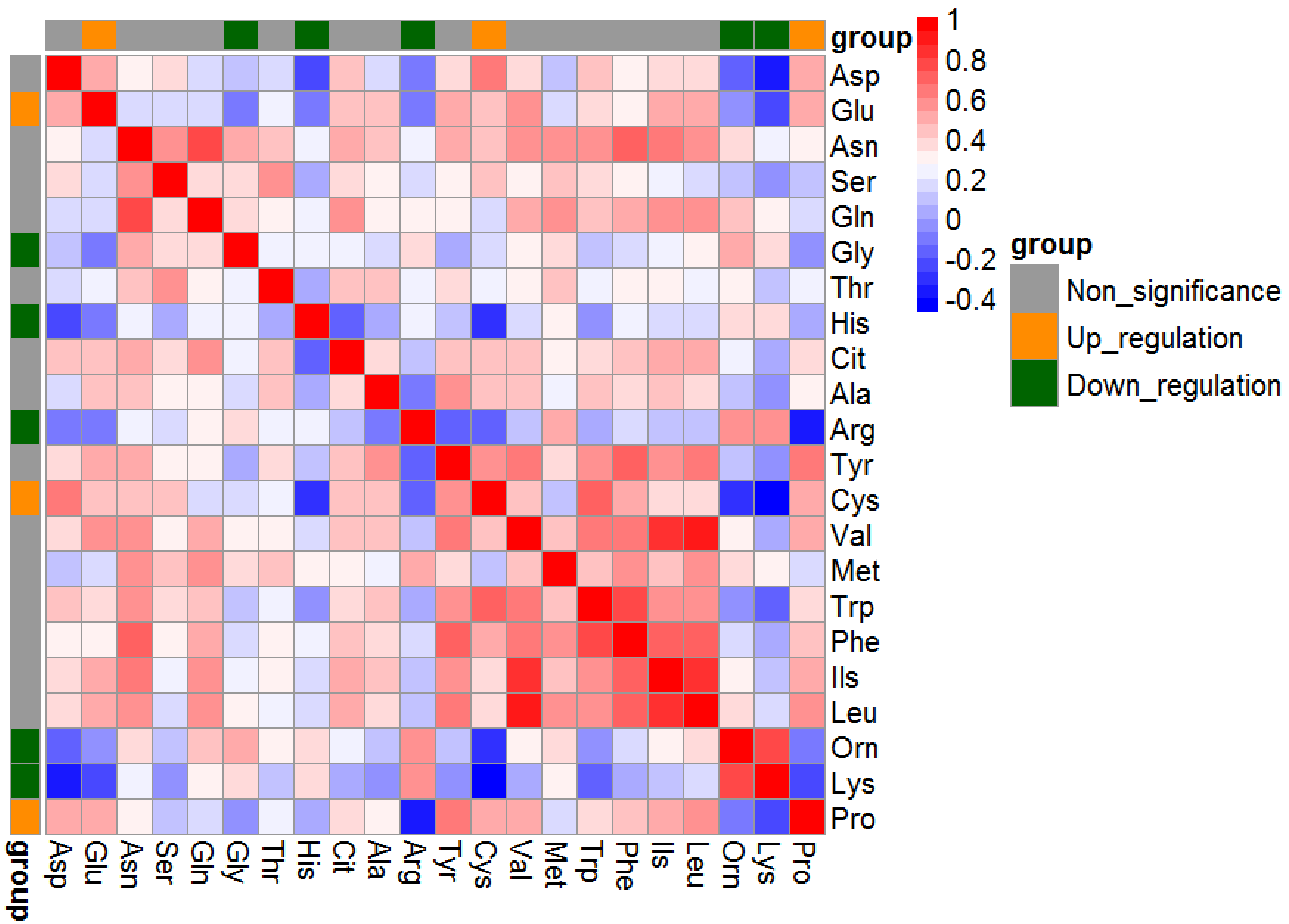
